# Occidiofungin, an actin binding antifungal with *in vivo* efficacy in a vulvovaginal candidiasis infection

**DOI:** 10.1101/368720

**Authors:** Akshaya Ravichandran, Mengxin Geng, Kenneth G. Hull, Daniel Romo, Shi-En Lu, Aaron Albee, Christopher Nutter, Donna M. Gordon, Mahmoud A. Ghannoum, Steve W. Lockless, Leif Smith

## Abstract

Current antifungal treatment options are plagued with rapidly increasing occurrence of resistance, high degree of toxicity and a limited spectrum of activity. The need to develop a novel antifungal with a unique target, wider spectrum of activity, and reduced toxicity to the host, is urgent. We have identified and characterized one such compound named occidiofungin that is produced by the soil bacterium *Burkholderia contaminans* MS14. This study identifies the primary cellular target of the antifungal, which was determined to be actin. Actin binding metabolites are generally characterized by their ability to inhibit polymerization or depolymerization of actin filaments, which presumably accounts for their severe toxicity. Occidiofungin, instead, has a subtler effect on actin dynamics that triggers apoptotic cell death. We were able to demonstrate the effectiveness of the antifungal in treating a vulvovaginal yeast infection in a murine model. This discovery puts occidiofungin in a unique class of actin-binding antifungal compounds with minimal reported toxicity to the host. The results of this study are important for the development of a novel class of antifungals that could fill the existing gap in treatment options for fungal infections.

**Author summary:** Widespread resistance to antifungal compounds currently in use has been alarming. Identification and development of a new class of antifungals with a novel cellular target is desperately needed. This study describes the assays carried out to determine the molecular target and evaluate efficacy of one such novel antifungal compound called occidiofungin. Occidiofungin modified with a functional alkyne group enabled affinity purification assays and localization studies in yeast. These studies led to the identification of the actin binding property of occidiofungin. Actin-binding by secondary metabolites often exhibit severe host toxicity, but this does not appear to be the case for occidiofungin. We have previously been able to administer occidiofungin to mice at concentrations in the range of 5 mg/kg without any serious complications. We were able to demonstrate the effectiveness of the antifungal in treating a vaginal fungal infection in a murine model. The results outlined in this manuscript establish that occidiofungin is an efficacious compound with a novel molecular target, putting it in a completely new class of antifungals.

## Introduction

Fungal infections caused by pathogens that are resistant to commonly used classes of antifungals are becoming increasingly prevalent. Recently, a CDC report described the spread of multi-drug resistant *Candida auris* causing systemic infections in hospitalized patients [1]. Furthermore, other species such as *Candida glabrata* and *Candida parapsilosis* have been reported to have gained resistance to routinely used azoles and echinocandins [2-4]. An example of a fungal infection that is rapidly developing resistance to currently available forms of treatment is vulvovaginal candidiasis (VVC). VVC will affect approximately 75% of all women and 5-10% of all women will develop recurrent VVC (RVVC) [5-7]. Approximately 90% of VVC is caused by *C. albicans*, while the remaining 10% is caused by *C. glabrata*, *Candida tropicalis*, *Candida parapsilosis,* and *Candida krusei*. These non-albicans VVC causing species are generally resistant to azole treatments, which are the most common treatment option for VVC [3, 8]. There have been no new therapeutic developments in decades for recurrent VVC. In the absence of well-established medical treatment standards, several ineffective methods for treating VVC have been prescribed, which generally include the use of a rigorous dosing regimen of antifungals followed by a long period of prophylactic dosing [5, 9, 10].

*Candida* species are an important cause of infection and mortality in all hospitalized patients[11]. Candidemia has a mortality rate of 30%–50% in cancer patients and is a major complicating factor for the successful treatment of cancer. Despite current antifungal drugs, invasive fungal infections are still a major cause of morbidity and mortality in transplant patient population[12, 13]. Reports suggest that candidal infection is the first and second most common infection in lung and heart transplant recipients, respectively[14-17]. In heart transplant recipients, candidal disease has been attributed to a mortality rate of 28%[18]. A rise in candidemia caused by non-albicans *Candida* spp. and an increase in azole resistance[19-23] is alarming; and supports the need for new antifungals. This problem is expected to be exacerbated by the presence of the multidrug resistant fungi *Candida auris* in hospitals. The increase in *C. auris* infections is expected to further reduce the positive therapeutic outcomes associated with currently approved antifungals [21]. Additionally, more appropriate antifungal treatment options may reduce the cost of treatment and mortality of patients.

Clinically approved antifungals primarily comprise members of the polyene, echinocandin, and azole family of compounds. The polyene antifungal amphotericin B was introduced in the 1950s and was the only antifungal available until the introduction of the azole class of antifungals in the 1980s. These two groups primarily target ergosterol production or bind to ergosterol, disrupting the fungal membrane. The echinocandins, the third group, are synthetically modified lipopeptides that originate from a natural cyclic peptide compound produced by fungi. This group selectively inhibits 1,3-β-glucan synthesis by functioning as a non-competitive inhibitor of 1,3-β-glucan synthase [24-27]. Widespread resistance and the ineffective spectrum of activity of this class of antifungals have been reported [28-32]. The prevalence of echinocandin- and azole-resistant fungal pathogens and the limited spectrum of activity of those compounds is one major issue contributing to the need for a new class of antifungals. Additionally, current antifungal treatments lead to abnormal liver and kidney function tests and have limitations with respect to their spectrum of activity and toxicities [33, 34]. Presumably, the identification of a novel class of antifungals with a broad spectrum of activity and a unique mechanism of action would mitigate the loss of life associated with the use of the current classes of antifungals. These limitations and toxicity problems have created an urgent need to identify antifungal compounds that have a novel mechanism of action [35].

Occidiofungin is a non-ribosomally synthesized glycolipopeptide produced by the soil bacterium *Burkholderia contaminans* MS14 [36]. It is a cyclic peptide with a base mass of 1200 Da (Fig 1). The bacterium produces a mixture of structural analogs of the base compound (occidiofungins A-D), however all analogs are composed of eight amino acids and a novel C18 fatty amino acid (NAA) containing a xylose sugar, and a 2,4-diaminobutyric acid (DABA). The structural analogs differ by an addition of oxygen to asparagine 1 (Asn1) forming a β-hydroxy asparagine 1 (BHN1) and by the addition of chlorine to β-hydroxy tyrosine 4 (BHY) forming 3-chloro β-hydroxy tyrosine 4 (chloro-BHY) [36]. Occidiofungin has a wide spectrum of activity against filamentous and non-filamentous fungi and minimal toxicity in an animal system [36, 37]. We have previously demonstrated that the mechanism of action of occidiofungin differs from the primary mode of action of the three common classes of antifungals [38, 39]. Briefly, there was no decrease in the activity of occidiofungin against *C. albicans* in the presence of 0.8 M sorbitol, an osmotic stabilizer, indicating that occidiofungin was not causing osmotic stress by cell wall or membrane disruption (the primary mechanism of action of azoles). Further, upregulation of Hog1p, which is an osmotic disruption indicator, was significantly lower than that seen for conditions known to induce the osmotic stress response pathway (e.g. 1 M NaCl). Mutants lacking Fks1p, an enzyme in the cell wall biosynthesis pathway, did not demonstrate increased resistance to occidiofungin. Disruption of the Fks1/Fks2 complex is the primary mechanism of action of echinocandins. Additionally, the introduction of vesicles containing ergosterol, the target of amphotericin, did not reduce the activity of occidiofungin unlike the case with amphotericin B. When observed under a microscope, occidiofungin-treated cells did not undergo lysis but appeared shrunken in size. Additional assays indicated that occidiofungin rapidly induces apoptosis in yeast cells at the minimum inhibitory concentration [39]. Interestingly, a critical threshold concentration of occidiofungin is required for its observed fungicidal activity. Occidiofungin has little impact on the growth rate of yeast at sub-inhibitory concentrations [38]. In addition, occidiofungin was seen to have potent inhibitory activity against *Pythium* species which lacks ergosterol in the membrane and against *Cryptococcus neoformans* which is resistant to echinocandins [36]. Preliminary toxicological analyses of occidiofungin using a murine model indicated that it was well tolerated at concentrations of 10 to 20 mg/kg [37]. Intravenous administration of occidiofungin to mice at a dose of 5 mg/kg was carried out with minimal induced toxicity. Blood chemistry analyses and histopathology performed on multiple organs showed a transient non-specific stress response with no damage to organ tissues [40]. Taken together, the data suggest that occidiofungin is a promising candidate for development as a clinically useful antifungal agent. This report describes studies to identify the molecular target of occidiofungin and determine its efficacy in a murine model of vulvovaginal candidiasis.

**Figure 1.**
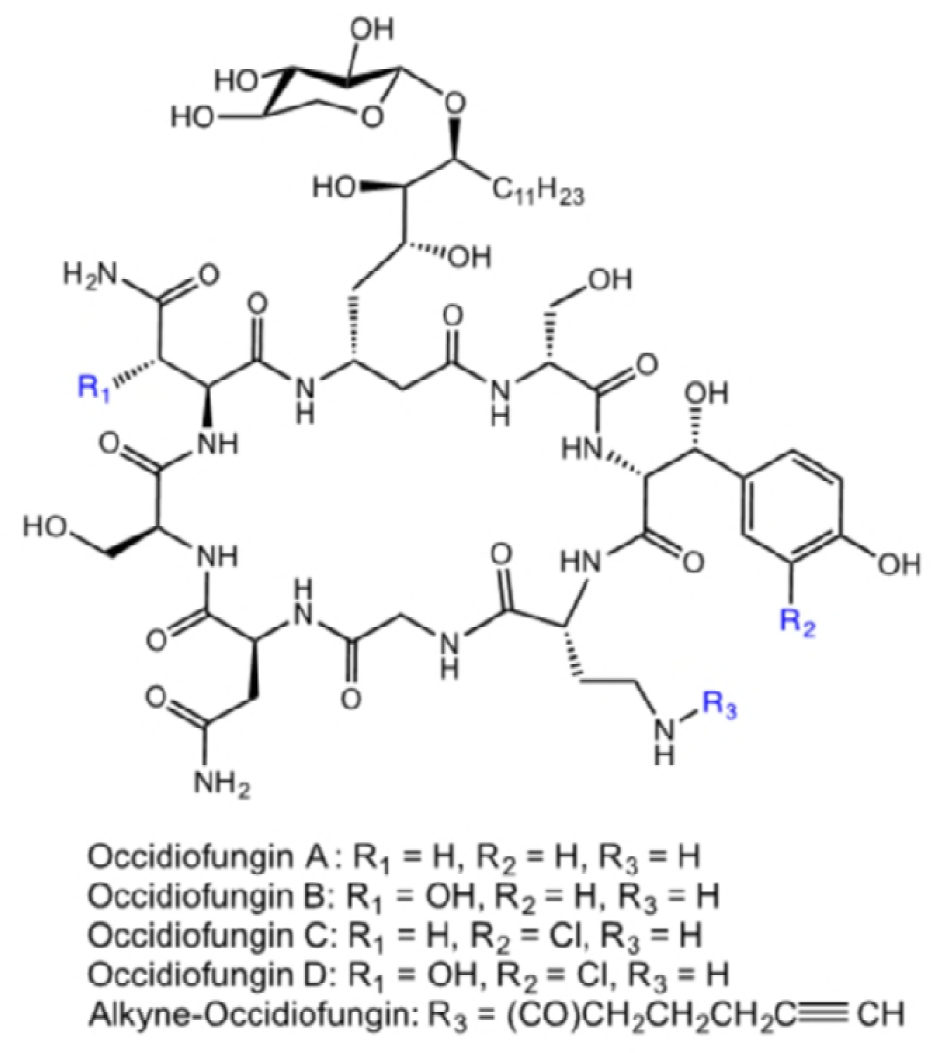
Covalent structure of occidiofungin A-D and alkyne-OF.

## Results

### Spectrum of activity of occidiofungin against clinically relevant fungi

Occidiofungin causes cell death in fungi through a mechanism of action that is distinct from the clinically used classes of antifungals [39]. Due to its unique mechanism of action, occidiofungin has sub-micromolar activity against azole and echinocandin resistant strains of fungi. Strains of *Candida albicans, Candida glabrata, and Candida parapsilosis* that were resistant to fluconazole and caspofungin were sensitive to occidiofungin (S1 Table). Furthermore, strains of *C. auris* were sensitive to occidiofungin at sub-micromolar concentrations. Non-albicans strains are believed to be the primary cause of recurrent vulvovaginal candidiasis. Strains of *Candida parapsilosis* and *C. neoformans* that were resistant to treatment with caspofungin were found to be susceptible to treatment with occidiofungin. Occidiofungin was also found to be active against *Aspergillus, Mucor, Fusarium,* and *Rhizopus* species. Several strains of the dermatophyte *Trichophyton* were also found to be susceptible to occidiofungin treatment, including azole and terbinafine resistant strains. A summary of the results, as reported in S1 Table, indicate that occidiofungin has activity against filamentous and non-filamentous fungi at sub-micromolar concentrations and has a broader spectrum of activity compared to other clinically available antifungals. Furthermore, sensitivity of fungal strains resistant to azoles and echinocandin class of antifungals support the notion that occidiofungin is functioning via a novel mechanism of action.

### Perturbation of actin-based functions following occidiofungin exposure

As a dimorphic fungus, *C. albicans* can grow as yeast or hyphae and the ability to switch between these forms is linked to the pathogenicity of the organism [41]. As most studies on occidiofungin efficacy have been carried out on *C. albicans* in their yeast form, the impact of the antifungal on morphological switching was tested. Incubation of *C. albicans* with sub-inhibitory concentration of occidiofungin was shown to block hyphae formation in cells that were induced to undergo morphological switching (Fig 2). Morphogenesis of *C. albicans* from yeast to filamentous forms has been shown to involve actin dynamics as treatment with cytochalasin A, latrunculin A, or the elimination of myosin I function prevent hyphae formation [42, 43]. In *C. albicans*, maintenance of the actin scaffold is also necessary for endocytosis, DNA segregation, and cell division [44, 45]. To determine whether occidiofungin impacts other cellular activities linked to actin dynamics, the effect of occidiofungin on endocytosis in fission yeast was evaluated by staining cells with FM-464 (Fig 3). Cells exposed to 0.5X MIC and 1X MIC demonstrated a concentration dependent reduction in stained endocytic vesicles. Actin has also been linked to the proper positioning of the mitotic spindle during cell division, and mutants that lack actin cables have been shown to accumulate multinucleated cells [46, 47]. Within thirty minutes of exposure, both *S. cerevisiae* and *C. albicans* cultures treated with a sub-inhibitory concentration of occidiofungin were found to accumulate a low percentage of binucleated cells indicative of a disruption or a delay in nuclear transit through the mother-daughter neck (S2 Table). To further characterize the role of actin in cellular response to occidiofungin, we analyzed haploid *S. cerevisiae* mutants deleted for genes linked to actin polymerization and depolymerization. Of the eighteen strains tested, only the Δ*tpm1* mutant showed altered sensitivity to occidiofungin, with the deletion mutant exhibiting a four-fold resistance to occidiofungin [S3 Table]. The observed increase in resistance to occidiofungin in the absence of the *tpm1* gene, which codes for the major isoform of tropomyosin, may be due to the mutant’s increased tolerance of cellular stressors (unpublished data) or a decrease in cellular growth rate [48]. A decrease in cellular growth has previously been linked to occidiofungin resistance [49]. The lack of an observed effect on occidiofungin activity with the vast majority of the major actin associated proteins suggests that they are not directly involved in the observed inhibitory activity of occidiofungin.

**Figure 2.**
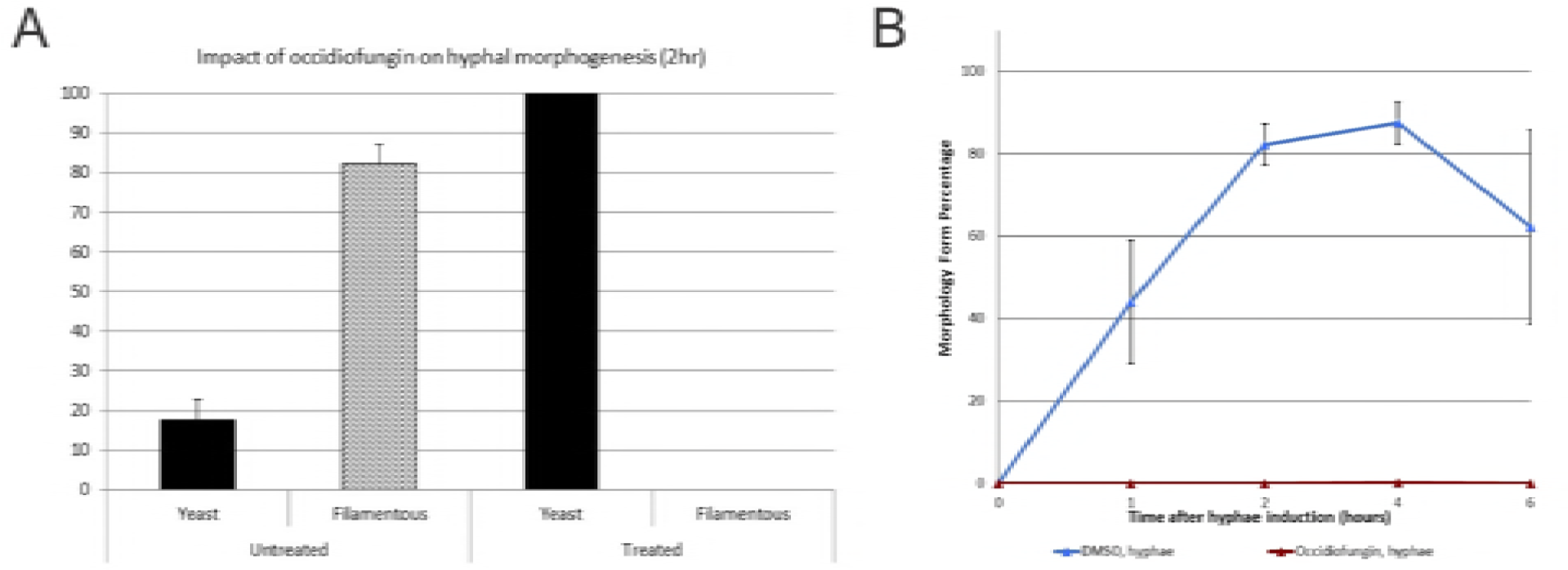
*Candida albicans* morphology under hyphae inducing conditions. (A) The resulting morphology was scored as either ‘yeast’ or ‘filamentous’ at two hours and the resulting percent given. The data is presented as the average with the standard deviation for over 200 cells from each treatment condition (n=3). (B) The resulting cell morphology was scored as either ‘yeast’ (open circles) or ‘filamentous’ (closed triangles) after 0, 1, 2, 4, and 6 hours at 37°C. The data is presented as the average with the standard deviation for over 200 cells from each treatment condition (n=3). DMSO treated samples are represented by blue lines; Occidiofungin treated samples represented by red lines.

**Figure 3.**
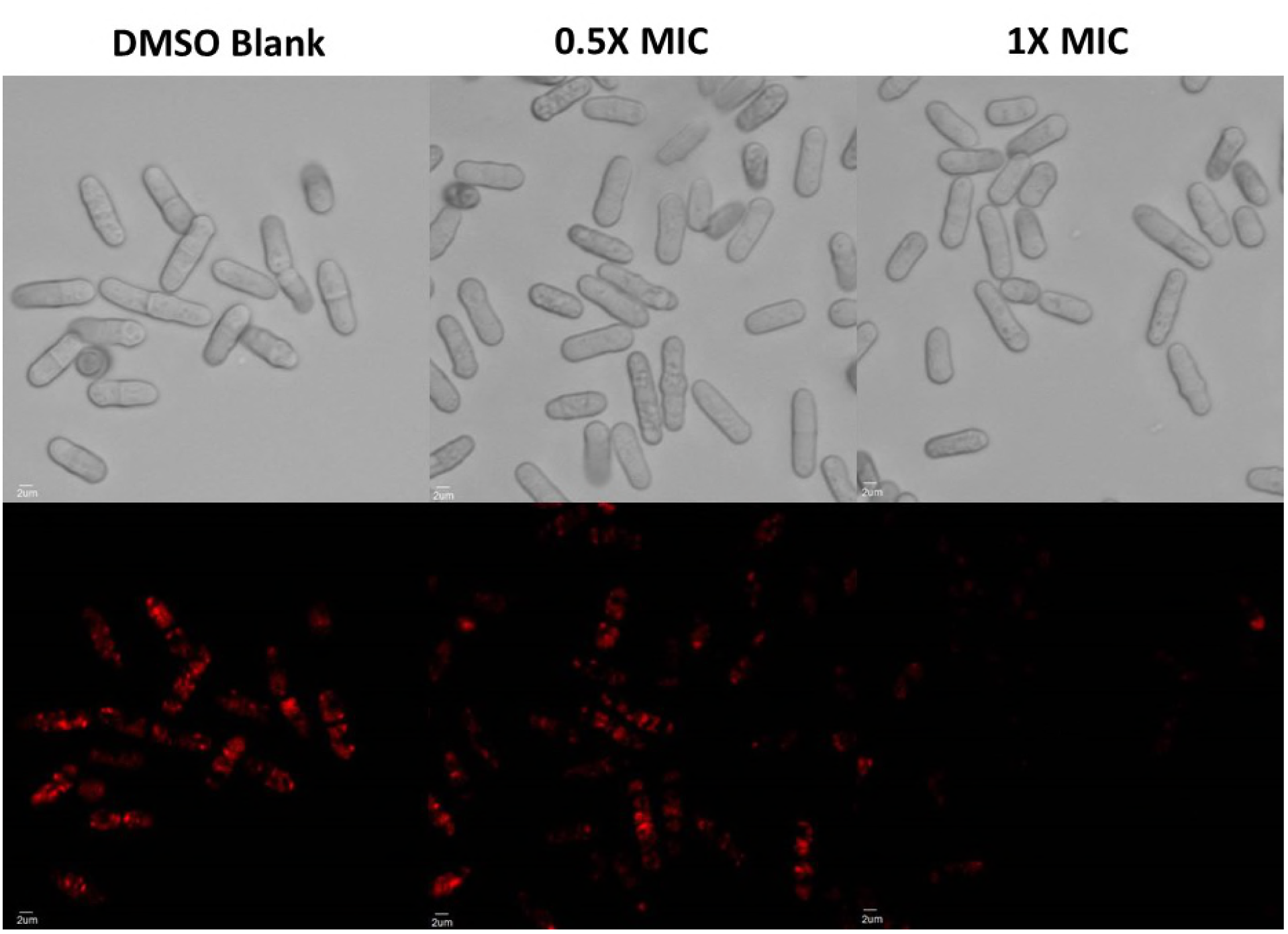
Effect of the native occidiofungin on endocytosis in fission yeast: DIC (top row) and fluorescence (bottom row) images of cells stained using FM-464 following treatment with sample blank (left column), 0.5× MIC of occidiofungin (middle column), and 1× MIC occidiofungin (last column). FM-464 dye uptake by endocytosis decreases in cells exposed to occidiofungin a dose dependent fashion.

### Alkyne derivatization of occidiofungin

In order to localize occidiofungin in yeast and to identify its cellular binding partners, methods to fluorescently label or add a functional purification tag to occidiofungin were needed. To this end, occidiofungin was chemically modified with a terminal alkyne through acylation of the free amino group of the diaminobutyric acid residue at position 5 (S1 Figure) for subsequent click chemistry (Sharpless-Hüisgen cycloaddition). Structural analysis of the derivatized product (S2 Figure and S3 Figure) revealed that occidiofungin (OF-B) and burkholdine (Bk-1215)[50], isolated by Schmidt, are likely identical products. The modified occidiofungin, alkyne-OF, had an eight-fold reduction in activity with the minimum inhibitory concentration of 1 and 0.5 μg/mL against *Saccharomyces cerevisiae* BY4741 and *Schizosaccharomyces pombe* 972 h-, respectively (S4 Table). To determine whether alkyne-OF still had the same apoptosis inducing bioactivity as the native occidiofungin, *S. cerevisiae* was treated with alkyne-OF and apoptotic assays such as TUNEL, reactive oxygen species (ROS) detection, and phosphatidylserine externalization assays were performed. Double stranded DNA breaks, the generation of ROS, and the externalization of phosphatidylserine were observed in the alkyne-OF treated cells, supporting the same mechanism of action (S4 Figure A-C). Although this alkyne modification moderately reduced the inhibitory activity of the compound, the functionalized derivative has the same apoptotic bioactivity and was therefore used to identify the fungal target.

### Identification of occidiofungin interacting proteins

Alkyne-OF was used in a pull-down assay to identify intracellular proteins that directly or indirectly interact with occidiofungin (Fig 4A). In brief, alkyne-OF was incubated with *S. cerevisiae* cells and the resulting cell lysates subsequently reacted with biotin-azide with occidiofungin-interacting proteins captured by passage over streptavidin beads. In the SDS PAGE gel, samples that included alkyne-OF had a more pronounced Coomassie blue stained band of captured proteins compared to control samples (Fig 4A). These bands were removed from the SDS PAGE gel for subsequent LC-MS/MS analysis. A silver stained gel containing samples electrophoresed to completion is provided to further show that the alkyne-OF was more efficient at capturing proteins in the pull-down assay. Data from multiple analyses using *S. pombe* 972h- and *S. cerevisiae* BY4741 were pooled. The resulting list of proteins obtained following LC-MS/MS analysis of excised bands was distilled as follows. Proteins that were observed in the two control samples, DMSO treated and native occidiofungin treated, were removed from consideration resulting in proteins that were exclusively found in the test sample captured with the alkyne-OF variant (S5 Table). The culled protein list was grouped based on gene ontology including cellular localization and/or molecular function. The resulting distribution is presented in Fig 4B. This analysis revealed that the majority of the proteins pulled down by alkyne-OF were actin or actin associated proteins (e.g. Pil1 and Cap1). In addition to actin-related proteins, proteins involved in vesicle transport and mannosylation were found associated with alkyne-OF. The remaining proteins were ribosomal and mitochondrial related proteins. The data indicates that occidiofungin plays a role in binding to actin since a majority of the proteins either directly influence actin dynamics (e.g. Arp2/3), are in close proximity to actin patches within the cell, or utilize actin filaments for their activity (e.g. Myo1).

**Figure 4.**
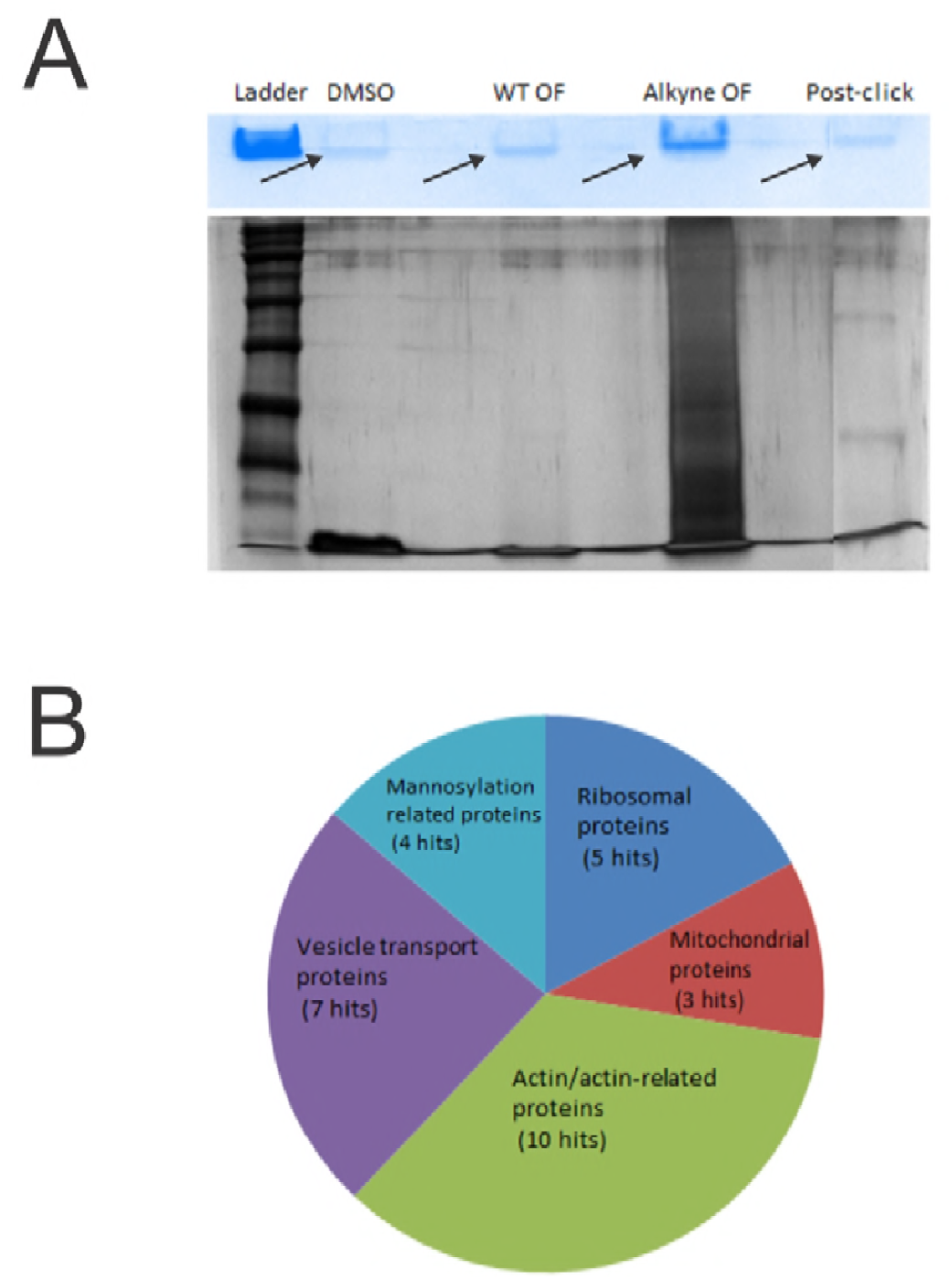
Determination of *in vivo* interaction of occidiofungin: A) Representative samples obtained following affinity purification of whole cell extracts run on 12% SDS PAGE gels and stained with Coomassie blue (top) and silver staining (bottom). The Coomassie stained gel was run only until the proteins entered the separating phase whereas the silver stained gel was allowed to run completely. The bands in the Coomassie stained gel (demarcated by the arrows) were removed and used for LC-MS/MS analysis to determine the proteins. Broad range (10-250kDa) ladder was used on both gels; B) Cellular distribution of the proteins obtained in the pull-down assay following LC-MS/MS analysis.

### In vitro analysis of the interaction of occidiofungin with purified actin

Typical assays for characterizing actin binding natural products are the *in vitro* F-actin polymerization and depolymerization experiments. However, the addition of occidiofungin was found to have no effect on the polymerization or depolymerization properties of F-actin (S5 Figure). Therefore, additional studies were required to confirm that actin was the biological target for occidiofungin. Biotinylation of alkyne-OF following incubation with F- or G-actin and streptavidin agarose beads was performed to determine whether occidiofungin directly associated with purified actin *in vitro*. F- or G-actin incubated with the wild type occidiofungin or DMSO were used as controls for potential non-specific interaction of actin with the agarose beads. The eluant from the biotinylated alkyne-OF had a single band at approximately 42 kDa, the expected size for actin. As shown in Fig 5A, the biotinylation of alkyne-OF was required for the co-purification of F- or G-actin with the streptavidin beads (Lane 5 and 8) as actin was not present in the control lanes that exposed actin to native OF or the carrier solvent DMSO (lanes 6, 7, 9, and 10). In this *in vitro* interaction assay, occidiofungin was shown to directly bind to F- or G-actin. To further support this observation, a dissociation constant of 1.0 ± 0.8 μM was determined from three independent ITC experiments using rabbit skeletal muscle G-actin (Fig 6). The ITC data also showed a 1:1 binding ratio for occidiofungin to G-actin. The ITC experiments were not adaptable to observe F-actin binding, so a co-sedimentation assay, which is commonly reported for identifying actin associated proteins, was performed [51, 52]. Phalloidin was used as a positive control in the assay (Fig 7). Phalloidin had an estimated dissociation constant (Kd) of 8 nM with a saturation of binding ratio of 0.6 phalloidin to one actin monomer. These values are corroborated by previous reports [53]. The biggest difference in binding to F-actin between occidiofungin and phalloidin is the number of molecules bound before saturation. Phalloidin saturated at approximately one molecule for every two actin monomers, whereas 24 molecules of occidiofungin were bound to each actin monomer before saturation. Even with the large number of bound occidiofungin to F-actin, the estimated Kd value was still 1.0 μM. It is important to note that the higher Kd value is attributed to a 50-fold increase in the amount of occidiofungin bound to actin compared to phalloidin.

**Figure 5.**
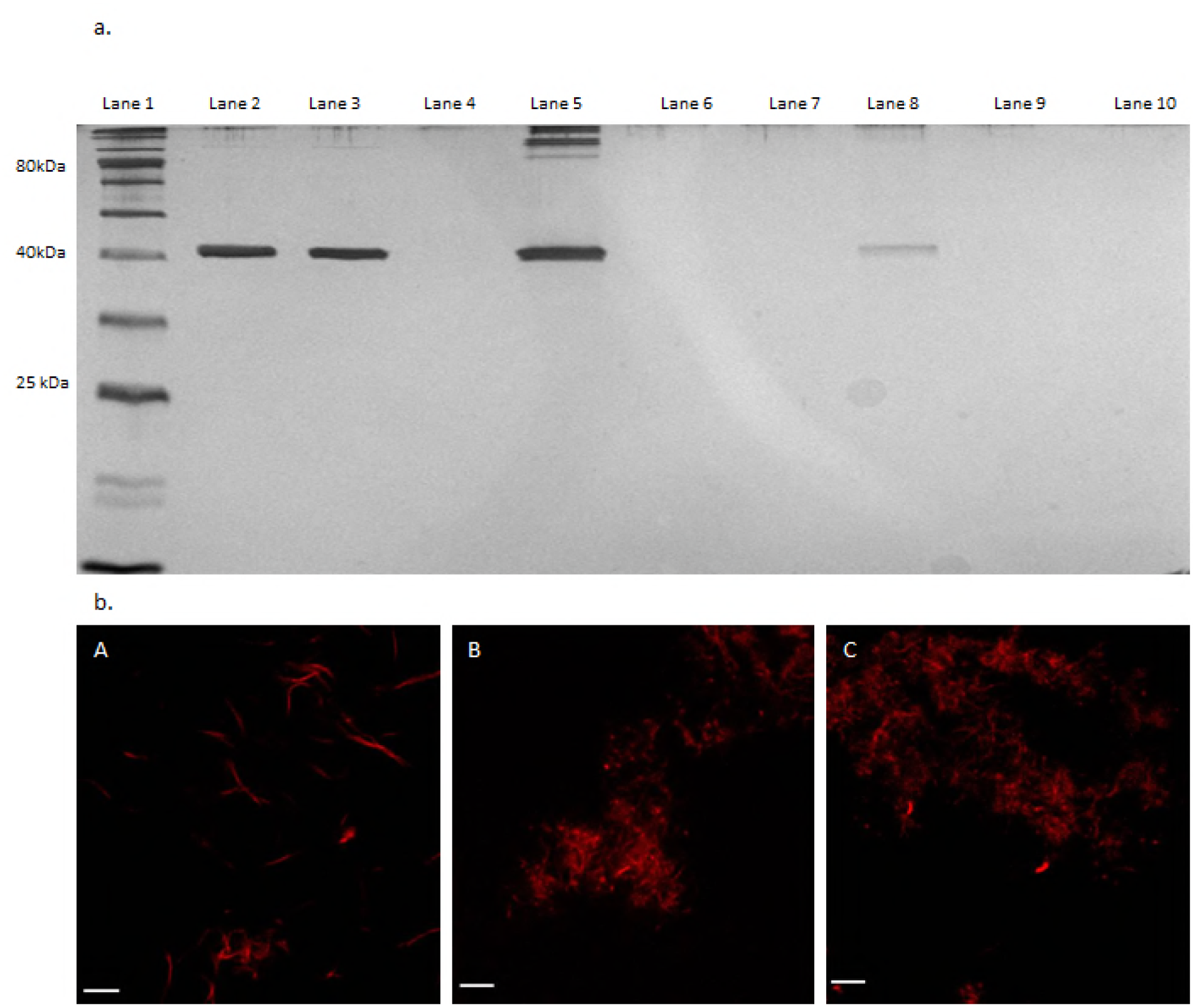
*In vitro* interaction of occidiofungin with F- and G-actin: a) Affinity pulldown of actin using alkyne-OF: Lane 1-Ladder, Lane 2-100 ng pure F-actin, Lane 3-100 ng pure G-actin, Lane 4-Empty, Lane 5-F-actin treated with alkyne-OF, Lane 6-F-actin treated with native occidiofungin, Lane 7-F-actin treated with DMSO, Lane 8-G-actin treated with alkyne-OF, Lane 9-G-actin treated with native occidiofungin, Lane 10-G-actin treated with DMSO; b) Fluorescence microscopy analysis of the effect of occidiofungin treatment on actin filaments visualized using fluorescently labeled phalloidin: A: actin filaments treated with solvent blank (DMSO), B: Actin:native occidiofungin (24 μg actin:4 μg native occidiofungin), C: Actin:native occidiofungin (24 μg actin:8 μg native occidiofungin). Scale bar represents 5μm.

**Fig 6.**
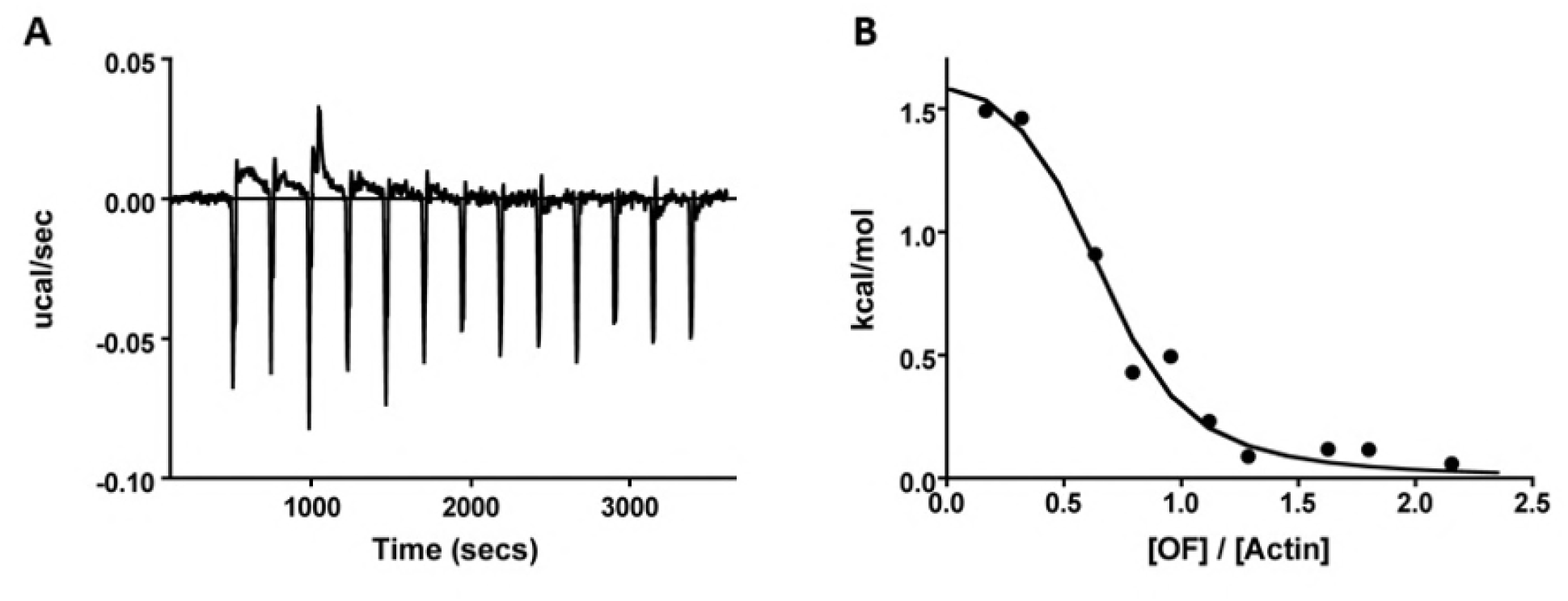
Isothermal titration calorimetry is used to measure occidiofungin (OF) binding to actin. A) Thermogram showing the heat exchange from equal injections of a solution containing OF into the ITC chamber containing actin. B) A representative binding isotherm of the integrated heat change from each injection shown in A is fit to a single-ligand binding model to yield an OF-actin dissociation constant. The mean and standard deviation from three independent experiments are K_D_ =1.0 ± 0.8 μM.

**Fig 7.**
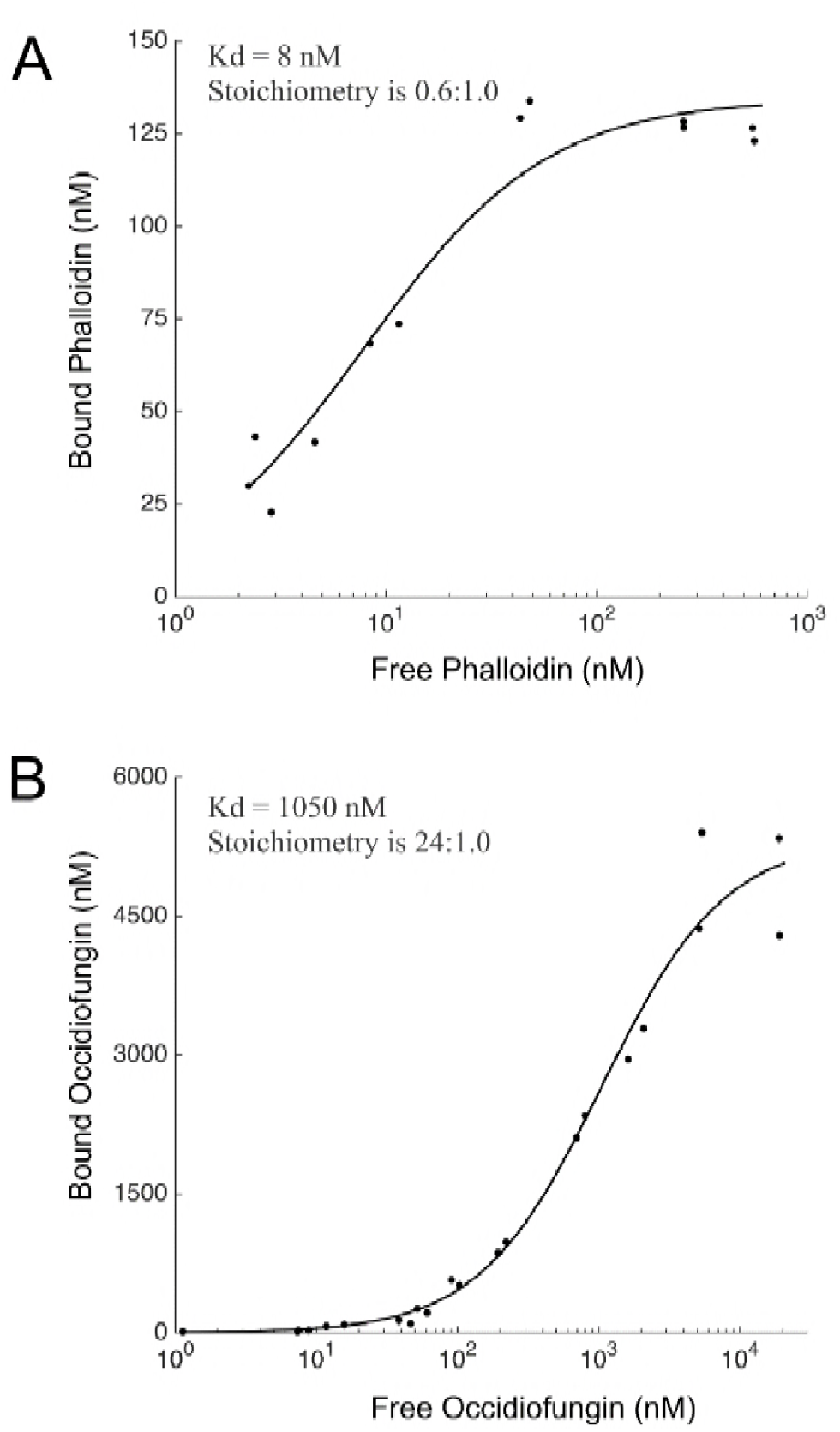
Co-sedimentation assay to demonstrate binding of occidiofungin to actin. A) Binding curve of phalloidin to actin (Kd = 8 nM and the stoichiometry (ligand: protein) is 0.6:1.0) and B) Binding curve of occidiofungin to actin (Kd = 1μM and the stoichiometry (ligand: protein) is 24:1). The graph is plotted between amount of free occidiofungin obtained in the supernatant of the co-sedimentation assay and the amount of bound occidiofungin obtained from the actin pellet. The data was fit to a standard Langmuir binding isotherm of the form: [X]bound = [X]*S/(Kd + [X]), where S is the maximal X bound, Kd is the dissociation constant and X is the concentration of free ligand.

Confocal microscopy using the fluorophore Acti-stain 670 phalloidin was carried out to visualize the impact of occidiofungin or alkyne-OF on actin filaments. Microscopic evaluation found that F-actin was still present but the morphology of F-actin changed in the presence of occidiofungin in a dose-dependent manner (Fig 5B). Using alkyne-OF labeled with azide functionalized Alexa Fluor 488 dye, occidiofungin interaction with F-actin was directly observed (S6 Figure). Similar to that shown using labeled phalloidin, F-actin also appeared to aggregate following exposure to occidiofungin. Fluorescence visualization of this interaction following treatment with alkyne-OF and native occidiofungin demonstrated a high degree of aggregation of the filaments which was not observed in the controls. Occidiofungin does not prevent polymerization or depolymerization of F-actin, but it does bind to F-actin causing it to aggregate. Given these unique observations for the bioactivity of occidiofungin, additional *in vivo* studies were conducted to verify that actin is its biological target.

### In vivo analysis of occidiofungin interactions with actin

*In vivo* visualization of the localization of occidiofungin was done in intact yeast cells. Cellular localization of F-actin is well characterized in *S. pombe* and *S. cerevisiae.* Time course analysis of *S. pombe* following alkyne-OF treatment and derivatization with azide Alexa-488 showed a specific pattern of localization of the compound (Fig 8A). Alkyne-OF was seen to have a faint pattern of staining at the polar tips at 10 minutes post treatment, which subsequently increased in intensity at 30 minutes post treatment. Strong fluorescence was observed at the polar ends of the cell and at the septum of dividing cells. A similar assay done using *S. cerevisiae* showed localization of alkyne-OF at the bud tips at the early time points and staining throughout the parent cell at later time points (Fig 8B). The unique pattern formed was observed to be a combination of striated and inclusion-like structures. In both yeast systems, when cells were pre-treated with native occidiofungin prior to treatment with alkyne-OF, the observed cellular localization patterns disappeared (Fig 8 A & B, panels D, E, and F) indicating that alkyne-OF and occidiofungin compete for the same cellular target. The vesicular pattern observed at the later time points of exposure is indicative of endocytic vesicles that are coated with actin being circulated through the cell [54, 55]. Actin patches in the cells of *S. pombe* are seen at the cell tips in growing cells and at the division septum in dividing cells. Actin patches recruited to the division septum interact with myosin to form the acto-myosin ring which is instrumental in cell division [56]. The time course analysis in both types of fungal cells show localization of occidiofungin to the regions where actin is known to be localized.

**Figure 8.**
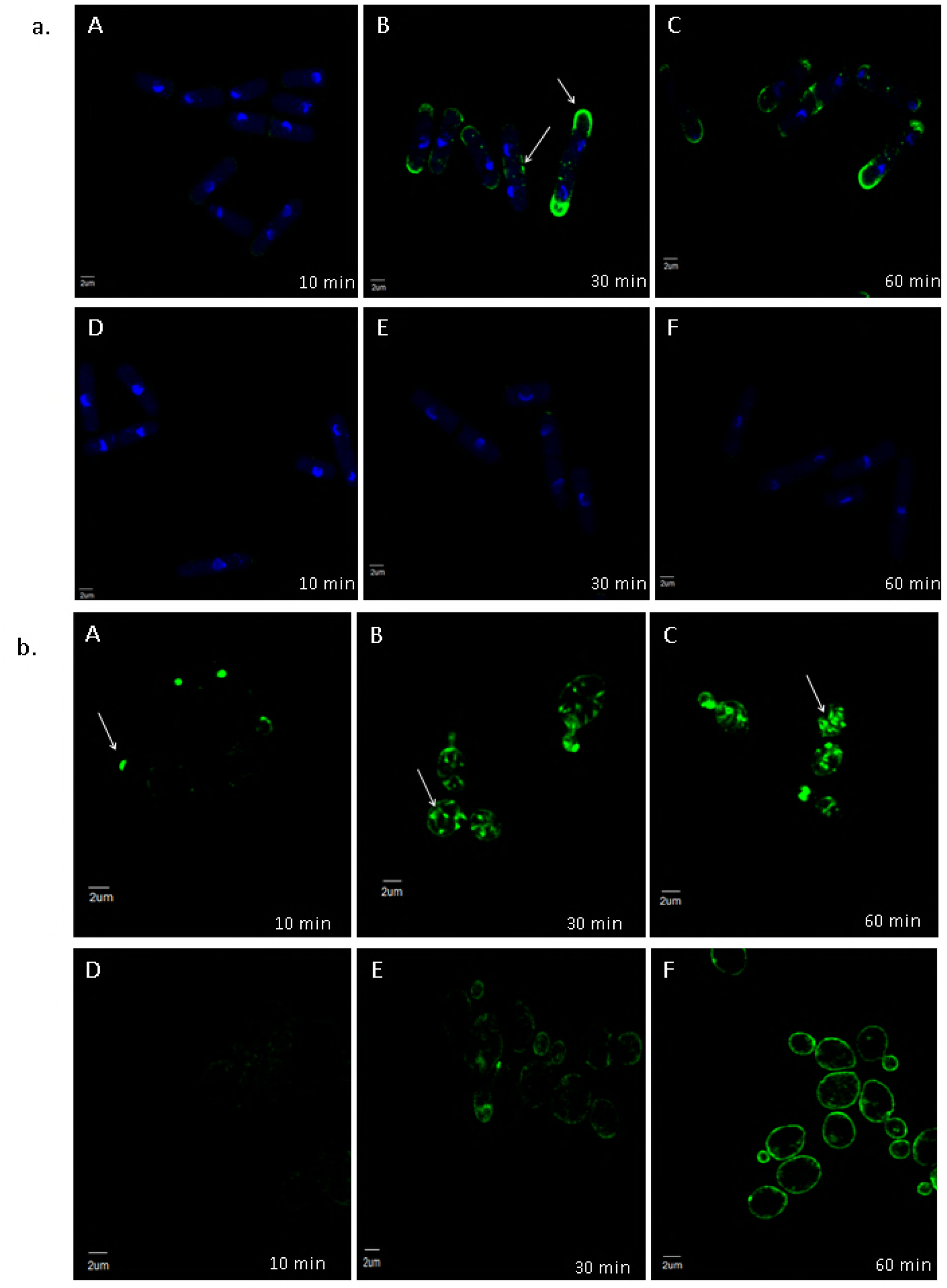
Competition assay of native occidiofungin and alkyne-OF. Time course analysis (A-C) of alkyne-OF distribution and the distribution of alkyne-OF with the competition of native occidiofungin (D-F) in a) *Schizosaccharomyces pombe* and b) *Saccharomyces cerevisiae*. Arrows indicate specific localization patterns of alkyne-OF observed in each cell at 10, 30, and 60 minutes. When pretreated with native occidiofungin, alkyne-OF does not bind or is restricted to cellular envelope in *Schizosaccharomyces pombe* and *Saccharomyces cerevisiae*, respectively.

To directly determine the effect of occidiofungin on actin organization *in vivo*, fluorescence microscopy was carried out on diploid cells of *S. cerevisiae* exposed to sub-inhibitory concentrations of occidiofungin. Within 30 minutes of exposure, an accumulation of actin patches and/or aggregates of F-actin were observed throughout the treated cells with a concomitant loss of actin cables (Fig 9A and 9B). Actin cables are formed by bundling F-actin. In Figure 8, the punctate structures in these cells are still likely filamentous actin, but occidiofungin appears to disrupt the organization of F-actin to form cables at sub-inhibitory concentrations.

**Figure 9.**
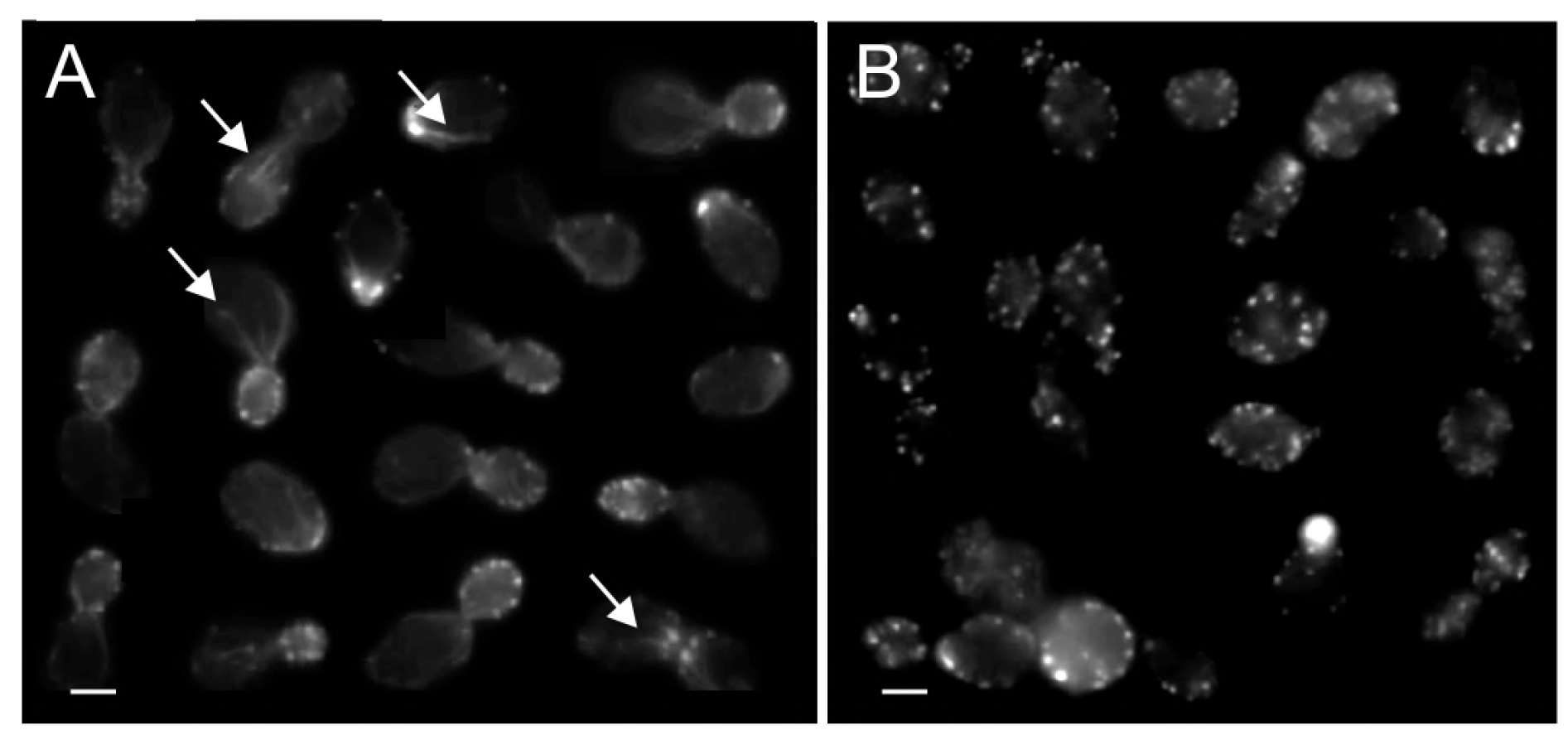
Effects of occidiofungin exposure on the integrity of actin cables in *S. cerevisiae* cells. A montage of cells processed for actin visualization using phalloidin-TRITC from: a) a culture exposed to solvent blank control (DMSO) where actin patches and cables are easily identifiable; and b) an occidiofungin treated culture (0.5X MIC; for 30 minutes) showing loss of actin cables and the accumulation of actin aggregates. Scale bars represent 2μm. The arrows are used to demarcate the presence of actin cables.

### Efficacy of occidiofungin in treating a murine model of vulvovaginal candidiasis

Six to eight-week-old BALB/c mice that were intravaginally infected with *C. albicans* were dosed once per day with occidiofungin for three days. The occidiofungin treated groups were compared to a vehicle control group. Three groups of six mice were treated with 100, 50, and 0 μg of occidiofungin suspended in 0.3% Noble agar. The occidiofungin treated groups reduced fungal load by more than two logs (Fig 10). The reduction in fungal load with both treatment groups was statistically significant from vehicle control (p<0.001). There was no statistically significant difference between the treated groups (p=0.33), suggesting that the lower limit of occidiofungin dosing was not achieved in the experiment. During the course of the study, the mice were examined for outward signs of distress or irritation. No behavioral changes including sluggishness, stretching, or reluctance to consume food was observed. Furthermore, no vaginal bleeding or swelling was observed following treatment.

**Figure 10.**
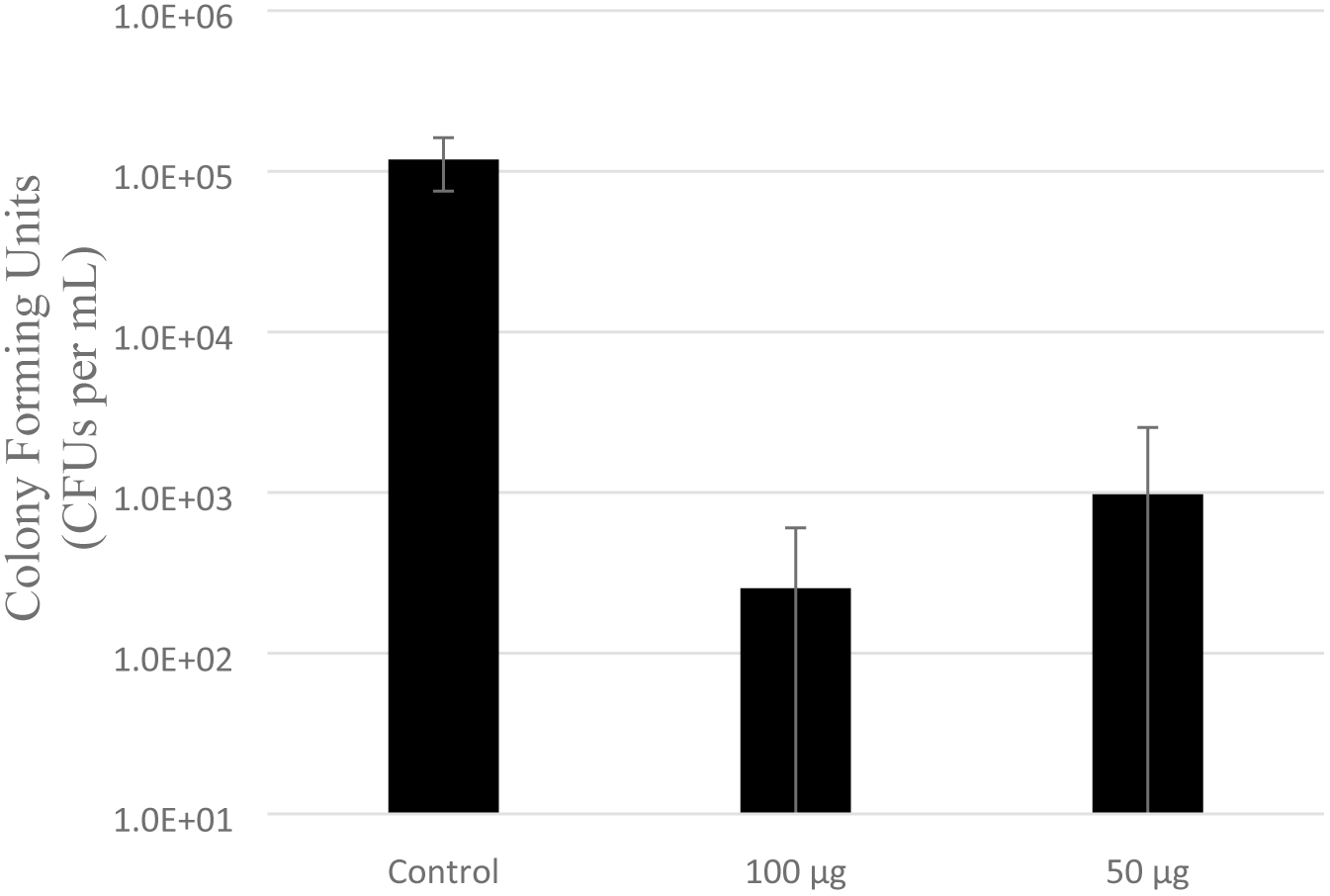
Efficacy of occidiofungin in treating murine vulvovaginal candidiasis. The graph demonstrates CFUs per ml of *Candida albicans* in the control group of mice compared to the groups treated intravaginally with different concentrations of occidiofungin in 0.3% noble agar. Error bars represent standard deviation. Statistical analyses indicate a significant difference between the control group and the treated groups (p<0.001) as indicated by the asterisk.

## Discussion

We have demonstrated that occidiofungin directly interacts with actin causing a disruption in normal cellular actin-based functions. *In vivo* data in yeast indicates that following treatment, the compound accumulates in areas rich in actin localization. In addition, disruption of cables throughout the fungal cells following exposure to sub-inhibitory concentrations of occidiofungin can be visualized. This interaction with actin can be observed *in vitro* via pulldown assays, ITC assays, and co-sedimentation studies. These studies establish direct interaction of occidiofungin with actin with a dissociation constant similar to other well-established actin binding proteins such as α-actinin, tropomyosin isoforms, and fimbrin [57-59]. From the binding studies and *in vitro* microscopy assays, it is evident that occidiofungin likely assembles into a large complex around F-actin. Based on the ITC study, occidiofungin and G-actin were bound at a 1:1 ratio. The propensity to form an occidiofungin complex appears to require the presence of F-actin. Formation of self-assembled complexes has previously been observed in several lipopeptide antibiotics [60]. It is likely that occidiofungin undergoes self-assembly, forming micellar structures at the concentrations tested in our assay. Additionally, cellular processes that rely on the maintenance of the actin cytoskeleton such as endocytosis, hyphae formation, and nuclear DNA positioning were shown to be disrupted with the addition of occidiofungin. We also demonstrate that occidiofungin is capable of treating a murine vulvovaginal candidiasis infection without any signs of toxicity. Furthermore, occidiofungin demonstrated efficacy at a concentration that is ten-fold lower than azole-based treatment methods [61].

Given that we have observed fungicidal activity as low as 100 nM concentrations, the number of bound occidiofungin to F-actin is likely to be a lot lower than 24:1 for its fungicidal activity. We hypothesize that occidiofungin binding to F-actin interferes with the binding of other actin associated proteins leading to disruption of cellular activities involving actin dynamics. Cables are necessary for a multitude of cellular functions including hyphal formation (disrupted following treatment as seen in Figure 2), endocytosis (reduced following treatment as seen in Figure 3) and proper positioning of the mitotic spindle during cell division (accumulation of multinucleated cells following treatment as seen in S2 Table). Studies aimed at understanding the events following the binding of occidiofungin to actin will need to be conducted to determine the exact connection for inducing apoptosis. Further, the localization pattern observed in the microscopy studies done in *S. cerevisiae* and *S. pombe* demonstrate the specificity of occidiofungin to F-actin. If the binding to actin were non-specific, actin would act as a retardant to the efficacy of occidiofungin due to the sheer number of occidiofungin molecules that are capable of binding a single F-actin monomer at a Kd value in the range of most actin associated proteins. The co-sedimentation assays and the localization studies indicate that the primary target of occidiofungin in the yeast cell is actin.

One of the challenges facing the development of antifungals is the fact that uptake of compounds into fungal cells does not occur as easily as it does in bacteria. Most fungal cells have a sturdy cell wall made of several glycoproteins that make up almost one-third of the dry weight of the cell. The efficiency of antifungals relies heavily upon being able to penetrate the cell envelope [62]. Occidiofungin has the advantage of being taken up by the fungal cell, as evidenced by the low MICs against several different types of fungi. Susceptibility to occidiofungin can be seen in pathogenic strains that are resistant to treatment with azoles and echinocandins (S1 Table). Additionally, occidiofungin has the advantage of inducing cell death in fungi via a mechanism that differs from the common classes of antifungals used to treat fungal infections.

Recent studies have shown that the dynamic nature of actin is necessary to maintain the cellular functions in which actin is involved such as endocytosis, mitochondrial transport, and growth [45]. A disturbance in the actin dynamics affects mitochondrial function [63, 64]. Previous reports suggest that stabilization and aggregation of actin leads to the induction of a Ras-cAMP-PKA pathway which causes mitochondrial destabilization and production of ROS [65, 66]. We have demonstrated that occidiofungin can directly interact with actin. Loss of actin cables following occidiofungin treatment can affect mitochondrial integrity, which in turn triggers a cascade of events leading to the release of reactive oxygen species. The release of ROS has been widely reported to be a precursor for the onset of apoptosis in yeast [67]. In addition, caspase dependent pathways have been theorized to be induced following aggregation of actin filaments in animal cells and it is possible that a similar pathway takes place involving Yca1p, the caspase found in yeast [68]. A newly formed bud contains a large pool of actin which coordinate the retrograde transport of vesicles along the actin cable into the mother cell. Actin nucleation is carried out by the Arp2/3 complex and a host of proteins including Cap1, Abp1 and Sac6 which are involved in the actin patch based transport of vesicles [45]. Fluorescence time course assays done on the cells of *S. cerevisiae* and *S. pombe* support this hypothesis. Furthermore, bud tips in *S. cerevisiae* are known to be rich in actin patches which are necessary to carry out cellular functions such as cell division and endocytosis [45]. Occidiofungin was observed to localize to these cellular areas in the *in vivo* microscopy experiments.

Natural products with *in vitro* antifungal properties that target the actin cytoskeleton have been previously reported [69]. One of the examples of an actin targeting antifungal is jasplakinolide, a compound that was isolated from sea sponges[70]. Although the compound had comparable activity against some *Candida* species when compared with miconazole, it had a limited spectrum of activity [70]. Furthermore, jasplakinolide was reported to be severely toxic in animal systems. It was reported to cause necrosis at the site of a subcutaneous injection at doses as low as 0.1 mg/kg and led to mortality at 1 mg/kg [71]. It is possible that its high level of toxicity is associated to the direct inhibition of actin depolymerization or its interaction with other cellular targets. A newly discovered actin binding antifungal is ginkbilobin [72]. This protein has been reported to induce programmed cell death following perturbation of the actin cytoskeleton and has a broader range of activity compared to jasplakinolide, however the effect of the compound in an animal system has not been reported [73]. Similarly, neosiphoniamolide A, a cyclodepsipeptide closely related to jasplakinolide, demonstrated a wide spectrum of antifungal activity. However, its toxicity in an animal system has not yet been reported [74]. Halichondramide, another antifungal compound isolated from a marine sponge, has been reported to be active against *Candida* species and *Trichophyton* species. Like jasplakinolide, it was severely toxic following a subcutaneous administration of 1.4 mg/kg of the compound in a murine system [75]. To date, the short list of actin binding antifungal compounds is limited by their toxicity in animals at doses that would be required to demonstrate an efficacious effect.

Occidiofungin, on the other hand, is well tolerated at 5 mg/kg or 20 mg/kg when administered intravenously or subcutaneously, respectively[37, 40]. This dose is much higher than the MIC of occidiofungin against several fungal pathogens. Furthermore, blood cells and blood chemistry following administration of occidiofungin show no serious signs of toxicity [40]. The difference in reported toxicity of occidiofungin to other actin binding compounds may be attributed to the inability of occidiofungin to disrupt polymerization or depolymerization of actin, different rates of uptake between yeast and animal cells, or the possible lack of off target binding that could lead to a toxic response. We have previously shown that occidiofungin is effective against several cancer cell lines [40]. The activity against these cell lines was almost a log more sensitive than the fibroblast cell line used as a control. It is believed that the growth rates of the cells are attributed to the differences in killing activity. This was later demonstrated with *S. cerevisiae, C. albicans and C. glabrata* strains of yeast grown in nutrient depleted conditions [49]. Further studies need to be carried out to examine the lack of severe toxicity of occidiofungin to an animal system. The low toxicity of occidiofungin combined with its wide spectrum of activity and demonstrable *in vivo* efficacy in treating a fungal infection is unprecedented in any actin binding antifungal compound.

Occidiofungin binds to actin and causes aggregation of the F-actin filaments *in vitro* which may lead to the accumulation of ROS and apoptosis *in vivo*. Demonstration of the effect of the compound in eliminating a common fungal infection *in vivo* supports the belief that occidiofungin could constitute a new class of clinically relevant antifungals. The activity of occidiofungin against echinocandin and azole resistant strains of pathogenic fungi leads us to consider that this compound would be a valuable addition to the existing antifungals for clinical treatment. Additional studies with occidiofungin may aid in furthering the understanding of the cellular events taking place that lead to the accumulation of ROS and apoptosis. A better understanding of entry and the events that lead to apoptosis following the binding of actin may lead to other potentially novel therapeutics. Further, future studies aimed at understanding how occidiofungin crosses the plasma membrane of fungi are warranted. Nevertheless, an actin-targeting antifungal that has a wide spectrum of activity against clinically pathogenic fungi and minimal toxicity in animal models could be the novel drug that is needed in the current antifungal arsenal to combat fungal infections.

## Methods

### Spectrum of activity of occidiofungin

Minimum inhibitory concentration (MIC) susceptibility testing was performed according to the CLSI M27-A3 and M38-A2 standards for the susceptibility testing of yeasts and filamentous fungi, respectively. Incubation temperature was 35°C and the inoculum size was 0.5 – 2.5 × 10^3^ colony-forming units (CFU)/mL and 0.4 − 5 × 10^4^ conidia/mL for yeasts and filamentous fungi, respectively. Inoculum concentration for dermatophytes was 1−3×10^3^ conidia/mL. RPMI was used throughout as the growth medium and *Cryptococcus* strains were tested in YNB. Occidiofungin MICs were recorded at 50% and 100% growth inhibition after 24 and 48 hours of incubation, with the exception of dermatophytes which were incubated for 96 hours. Fluconazole MICs against *Candida* strains were recorded at 50% inhibition after 24 hours and against *Cryptococcus* strains after 72 hours. Voriconazole MICs were recorded at 100% inhibition after 24 hours for zygomycetes and after 48 hours for *Fusarium* and *Aspergillus* strains. Voriconazole MICs were recorded at 80% inhibition after 96 hours of incubation for dermatophytes. *S. cerevisiae* deletion mutants were obtained from the commercially available BY4741 deletion library (Thermo Scientific). Susceptibility testing was carried out on inoculum size of 0.5−1×10^4^ cells/ml in YPD media at 30°C and MICs recorded after 48 and 60 hours.

### Hyphal induction

*Candida albicans* strain ATCC 66027 was grown in YPD at 30°C for 48 hours to reach saturation with a density of 17-19 OD_600_/ml. Cells were diluted into fresh Spider media (1% nutrient broth, 1% mannitol, 0.2% K_2_HPO_4_) to obtain 0.05 OD_600_/ml (~0.5−1×10^6^ cells/ml) and incubated at 37°C with shaking to induce hyphae formation[76]. Immediately prior to 37°C incubation, occidiofungin (1μg/ml; 0.5X MIC) or an equal volume of DMSO was added to cultures. Aliquots were removed at 0, 1, 2, 4, and 6 hour time points, fixed in 3.7% formaldehyde for later analysis of cell morphology by light microscopy. Over 200 cells were examined for each time point and cells were scored as having either a yeast or filamentous form with cells showing any outgrowth as being filamentous.

### DNA segregation

A mid log culture of *S. cerevisiae* (BY4743; diploid) and *C. albicans* (ATCC 66027) were diluted into fresh YPD to obtain an OD_600_/ml of 0.095 (~1-1.5×10^6^ cells/ml) and 0.05 (0.5-0.8×10^6^ cells/ml), respectively. DMSO or occidiofungin (1μg/ml; 0.5X MIC) was added and cultures were placed at 30°C with shaking. Samples were removed at 0.5, 1, and 2 hours and cells fixed for 1.5 hours at room temperature with the addition of formaldehyde to 3.7%. Cells were washed in PBS and permeabilized with the addition of an equal volume of PBS containing 0.2% TritonX-100 for 30 minutes at room temperature. Cells were isolated by centrifugation, washed in PBS, resuspended in a minimal volume of PBS, added to concanavalinA treated glass slide, and overlaid with VectaShield plus DAPI. Images were viewed using a Nikon 50i fluorescence microscope with a 100X oil immersion objective and DAPI filter. Cells were scored based on bud morphology and DNA location.

### FM-464 uptake assay for endocytosis

Three 1 mL aliquots of *S. pombe* at a density of OD_600_ 0.6 to 0.8 were treated with 0.01% DMSO, 0.5 μg/mL (0.5X MIC), or 1 μg/mL (1X MIC) of native occidiofungin for 30 minutes at 30°C. Cells were isolated by centrifugation at 21,000 g for 2 minutes, washed thrice with PBS and resuspended in YPD containing 8 mM FM-464 (ThermoFisher Scientific). Cells were incubated in the presence of the dye for 60 minutes at 30°C, followed by two washes with PBS and then added to a microscope slide for visualization. Images were obtained using an Olympus FV1000 confocal microscope with a 40x/0.9 dry objective.

### Derivatization of occidiofungin

Occidiofungin was purified from a liquid culture of *Burkholderia contaminans* MS14 as previously described [36]. Pure occidiofungin was aliquoted into 100 μg fractions and stored in lyophilized form at 4°C until use. Addition of an alkyne reactive group to the primary amine on occidiofungin was performed initially at the Texas A&M Natural Products LINCHPIN Laboratory at Texas A&M University and subsequently at the CPRIT Synthesis and Drug-Lead Discovery Laboratory at Baylor University.

The alkyne derivatization of occidiofungin was carried out under nitrogen atmosphere in a round bottom flask (S1 Figure) using HPLC grade reagents. A solution of triethylamine and acetonitrile was prepared by dissolving 10 μL of trethylamine in 11.7 mL of acetonitrile. A stock solution of acetonitrile and reagents was prepared by adding 2,5-dioxopyrrolidin-1-yl hex-5-ynoate (1.40 mg, 6.69 μmole, 8.2 equiv.) to 1.08 mL of the triethylamine and acetonitrile solution prepared above. Occidiofungin (1 mg, 0.82 μmole, 1 equiv.) was added to the stock solution of acetonitrile and reagents (400 μL) containing triethylamine (0.34 μL, 2.46 μmole, 3 equiv.) and 2,5-dioxopyrrolidin-1-yl hex-5-ynoate (51.6 μg, 2.46 μmole, 3 equiv.). Occidiofungin was added while stirring, with water (400 μL) added to the mixture as a co-solvent. The resultant mixture was stirred for three days at 22°C. Purification of alkyne-OF was performed on an Agilent 1200 series semi-prep HPLC (gradient 5% acetonitrile/water to 95% acetonitrile/water over 20 minutes) using a Phenomenex Gemini 5μ C18 110A (100 × 21.2 mm) reversed-phase column. For analytical analysis, a phenomenex Gemini 3μ C18 110A (150 × 4.6 mm) reversed-phase column was used on an Agilent 1200 series analytical HPLC system. Derivatized occidiofungin was also purified using a 4.6- by 250-mm C18 column (Grace-Vydac; catalog no. 201TP54) on a Bio-Rad BioLogic F10 Duo Flow with Quad Tec UV-Vis detector system using a similar gradient described above. Analytical HPLC analysis of the reaction mixture indicated complete consumption of the starting material and a new peak in the chromatogram (OF RT = 13.6 min; alkyne-OF RT = 15.0 min). The crude mixture was lyophilized and the crude powder, solubilized in 35% acetonitrile containing 0.1% trifluoracetic acid, was purified on a semi-prep HPLC (alkyne-OF RT = 13.0 minutes) column to yield alkyne-OF (0.76 mg, 70%).

NMR data were collected using a Bruker Ascend 600 (^1^H 600 MHz and ^13^C 150 MHz) NMR spectrometer equipped with a 5 mm cryoprobe. The NMR data for ^1^H NMR chemical shifts are reported as δ values in ppm relative to residual HOD (3.30 ppm), coupling constants (*J*) are reported in Hertz (Hz), and multiplicity follows convention. Unless otherwise indicated, dimethylsulfoxide-d6 (DMSO-d6) served as an internal standard (40.5 ppm) for the ^13^C spectra. NMR data for alkyne-OF A-D is shown in S2 Figure (A-F) and the assignment of the alkyne subunit to diaminobutyric acid (DABA5) is shown in S3 Figure (A-H). The assignment of the alkyne subunit has led to a correction in the previously reported configuration of DABA5^28^ and occidiofungin B is likely identical in structure to Bk-1215[50] isolated by Schmidt et al. Assignment of the High-resolution mass spectra (HRMS) were run on a Thermo LTQ Orbitrap mass spectrometer with ESI direct infusion (alkyne-occidiofungin B HRMS (ESI^+^): Calcd. For C_58_H_92_N_11_O_23_ ([M+H]^+^), 1310.6360. Found: 1310.6361).

### Actin polymerization and depolymerization assays

The effect of unmodified occidiofungin on actin polymerization and depolymerization was measured using the Actin Polymerization Biochem Kit (fluorescence format): rabbit skeletal muscle actin from Cytoskeleton Inc. (Catalog no.: BK003). Occidiofungin was brought up in 1.5% β-cyclodextrin in PBS (pH 7.5) at a concentration of 1 μg/μL and 20 μL of this solution was used per well. The same buffer without occidiofungin was used for buffer control. General Actin buffer was supplemented with ATP to 0.2mM prior to the start of the experiment as instructed. G-actin and F-actin stocks were prepared as described in the kit to achieve stock concentrations of 0.4 mg/mL and 1 mg/mL, respectively. Polymerization and depolymerization assays were then carried out as per the manufacturer’s instructions.

### Alkyne-OF functionality testing

Stock solutions of unmodified occidiofungin and alkyne derivatized occidiofungin were made in DMSO at a concentration of 1mg/mL. These stock solutions were utilized in all the assays described in this manuscript. The activity of the purified alkyne-OF was compared to the native compound using the CLSI M27-A3 method of determination of the minimum inhibitory concentration (MIC) against *Saccharomyces cerevisiae* BY4741 and *Schizosaccharomyces pombe* 972h (received from Dr. Susan Forsburg, Department of Biological Sciences, University of Southern California). Additionally, activity of the alkyne derivatized occidiofungin was tested against a higher density (OD_600_/mL 0.6 to 0.8) of cells for both yeast. Assays for TUNEL (APO™-BrdU TUNEL Assay Kit, LifeTechnologies), phosphatidylserine externalization (Annexin-V-Fluos staining kit, Roche), and ROS detection (Dihydrorhodamine 123, Sigma) were carried out as previously described[39].

### Affinity purification of alkyne-OF associated proteins

One mL of cells from an overnight culture of *S. pombe* grown to an OD_600_ of 0.6-0.8. was incubated with 8 μg/mL of alkyne-OF for 30 minutes at 30°C. Controls included cells treated with DMSO and native occidiofungin. An additional sample using cells that were lysed prior to alkyne-OF treatment was used for comparison (Figure 2A, Lane Post Click). Following exposure, cells were isolated by centrifugation, washed once with PBS and then lysed by probe sonication (30 second sonication followed by 30 second on ice; cycle repeated 5 times). Insoluble material was removed by centrifugation at 16,000 g for 10 minutes and the supernatant used in a Click-it protein reaction (Life Technologies) with the alkyne-OF reacted with azide-biotin, as per the manufacturer’s instructions. The reaction was allowed to proceed for 90 minutes at 37°C with shaking. Unreacted reagents were removed by passage through a Microcon 10 kDa cutoff filter (Sigma Aldrich) and the concentrated proteins solubilized in 100 μL of 100 mM Tris HCl (pH 7.5). Biotinlyated proteins were selected using streptavidin agarose beads (ThermoFisher Scientific) with incubation at 37°C for 90 minutes. Beads were washed with 10 mL of 100 mM Tris HCl (pH 7.5) and bound proteins extracted with boiling in 50 μL of 1X SDS sample loading buffer for 15 minutes. The resulting protein sample was loaded onto a 12% SDS gel and electrophoresis was carried out until the bromophenol blue dye was ~1cm into the separating gel. The resulting band was excised, trypsin digested by the Protein Chemistry Laboratory (Texas A&M University), and the subsequent LC-MS/MS analysis performed by the Mass Spectrometry Laboratory at the University of Texas Health Science Center (San Antonio). Results were analyzed using the Scaffold software.

### Intracellular localization of alkyne-OF

Colonies from a freshly streaked plate were used to inoculate an overnight culture of *S. pombe* or *S. cerevisiae.* 1 mL of cells from a culture with an OD_600_ of 0.6 to 0.8 was incubated with MIC quantities of alkyne-OF at 30°C with 200 μL samples removed at 10, 30, and 60 minute post incubation. Cells were isolated by centrifugation, washed once in phosphate buffered saline (PBS), and fixed for 15 minutes at room temperature with the addition of formaldehyde to 3.7% (in PBS). Cells were permeabilized at room temperature for 20 minutes with the addition of 0.5% TritonX-100 (in PBS). Cells were washed twice with 1 mL of PBS each. Click reaction with azide derivatized Alexa-488 was carried out according to the manufacturer’s protocol (Click-iT EdU Imaging kit, ThermoFisher Scientific). Cells were washed with PBS and added to a microscope slide for visualization using an Olympus FV1000 confocal microscope with a 100x/1.4 oil immersion objective and 40x/0.9 dry objective. A competition assay was carried out by pre-treating cells with an MIC amount (0.5 μg/mL) of the native occidiofungin followed by treatment with alkyne-OF.

### In vitro actin binding

Purified rabbit skeletal muscle filamentous actin (Catalog: AKF99) and G-actin (Catalog: AKL95) was purchased from Cytoskeleton Inc. Actin was reconstituted in Milli-Q water to achieve a stock concentration of 0.4 mg/mL in a buffer that consisted of 5 mM Tris-HCl (pH 8.0), 0.2 mM CaCl_2_, 0.2 mM ATP, 2 mM MgCl_2_, and 5% (w/v) sucrose as directed by the supplier. This solution was stored at −80°C in 50 μL aliquots until use. Immediately before use, each aliquot was thawed by placing in a 37°C water bath for 5 minutes followed by room temperature. For all studies, 24 μg of F- or G-actin was used with 8 μg alkyne-OF.

For biochemical-based studies, click chemistry was performed on this mixture to react the alkyne-OF with azide-biotin for 90 minutes as described (Click-iT protein reaction buffer kit, ThermoFisher Scientific). Unreacted reagents were removed by centrifugation (20 minutes at 15,000× g) through a 10 kDa cutoff filter. Proteins retained in the filter chamber were solubilized in 200 μL of 100 mM Tris-HCl (pH 7.5) and reacted with streptavidin beads as described above. The beads were washed multiple times using 100 mM Tris HCl (pH 7.5) and bound proteins eluted by boiling in 50 μl of SDS sample loading buffer. The sample was electrophoresed through a 12% SDS gel and protein bands visualized by silver staining according to the manufacturer’s protocol (Pierce Silver stain kit, ThermoFisher Scientific). F- and G-Actin treated with DMSO and native occidiofungin were used as controls.

For microscopy-based studies, click chemistry was performed to react the alkyne-OF with functionalized Alexa Fluor 488 according to the manufacturer’s instructions (Click-iT EdU Imaging kit, ThermoFisher Scientific). Unbound dye was removed by overnight dialysis at 4°C against actin polymerization buffer using a 1 kDa cutoff membrane (Catalog no.: BSA02, Cytoskeleton Inc.). Actin filaments were removed, added to a slide and analyzed using an Olympus FV1000 confocal microscope 40x/0.90 dry objective and a 100x/1.4 oil immersion objective. As a control, 140 nM Acti-stain 670 phalloidin (Cytoskeleton Inc.) was used to visualize actin filaments as per manufacturer’s instructions.

F-actin filaments were reacted with native occidiofungin for 15 minutes at room temperature at molar ratios of 1:10 (24 μg actin:8 μg native occidiofungin) and 1:5 (24 μg actin:4 μg native occidiofungin). The respective mixtures were then stained with 140 nM Acti-stain 670 phalloidin for an additional 15 minutes at room temperature. Stained filaments were added to a glass slide and observed on an Olympus FV1000 confocal microscope using a 100x/1.4 oil immersion objective. Actin filaments treated with DMSO (solvent blank negative control) were stained and observed for comparison.

### Isothermal Titration Calorimetry

Rabbit skeletal muscle actin (Cytoskeleton, Inc.) was re-constituted as described above. Aliquots of actin were thawed and incubated for 1 hour in ITC buffer which contained General Actin Buffer (Cytoskeleton, Inc.: 5 mM Tris-HCl, pH 8.0, 0.2 mM CaCl_2_) plus 0.5 mM DTT, 0.2 mM ATP and 5% DMSO. Lyophilized OF was re-suspended in ITC buffer to yield a 300μM concentration. The ITC chamber was loaded with 30 μM G-actin and the syringe with 300 μM OF. A MicroCal iTC200 (Malvern Instruments, Spectris plc) was used to obtain OF binding to actin at 25°C. A 0.3 μL injection was followed by 13 injections of 3 μL while stirring at 1,000 rpm. The integrated heat from each injection was fit to a one-site binding model using Origin.

### Co-sedimentation studies

Reaction buffer for F-actin binding study consisted of 5 mM Tris-HCl (pH 8.0), 0.2 mM ATP, and 2 mM MgCl_2_. Samples were prepared in 100 μL volume and the final concentration of F-actin was 200 nM. Occidiofungin was added to the reaction at concentrations from 25 to 25600 nM. Phalloidin was added to the reaction at concentrations from 25 to 800 nM. Samples were incubated at room temperature for 30 minutes in thick-wall polycarbonate tubes (#349622, Beckman Coulter, CA). After incubation, samples were centrifuged at 100,000 rpm, 4°C, for 20 minutes to pellet F-actin (Beckman TL-100 ultracentrifuge). Both supernatant and pellet were extracted with 200 μL solvent consisting of a combination of 70% acetonitrile (ACN) containing 0.1% trifluoroacetic acid (TFA) and 30% methanol containing 0.4% formic acid. The extracted sample was centrifuged at 15,000 g for 10 minutes, after which the supernatant was transferred to a clean centrifuge tube before being freeze dried in a SpeedVac (Labconco, Cat#7810010). The dried sample was brought up to 100 μL with 50% ACN with 0.1%TFA containing 1 μg/mL concentration of the internal standard and analyzed on LC-MS. An analog of occidiofungin served as the internal standard of native occidiofungin, while native occidiofungin served as the internal standard of phalloidin. All experiments were done in duplicate. An Agilent 1200 front end chromatography system and a TSQ Quantum™ Access Triple Quadrupole Mass Spectrometer were used to analyze the samples. Following a 10 μL injection, samples from binding study with occidiofungin were separated using a 15-minute water/ACN (containing 0.2% formic acid) gradient starting from 95% to 40% water on a C18 column (SinoChrom ODS-BP 5μm, 2.1 mm × 50 mm). The mass spectrometer was operated in positive mode and operated using a protocol optimized for phalloidin and occidiofungin. Briefly, occidiofungin was monitored in SRM mode with a scan width (m/z) of 0.3 and a collision energy of 31 eV, while phalloidin was monitored in SIM mode with a scan width of 0.7. The parent and product mass of native occidiofungin were 1216.7 and 1084.7 Da, respectively. The center mass of phalloidin was 789.87 Da. Area of each compound was measured through manual integration using Xcalibur™ Software (Thermo Fisher Scientific). The standard curves were generated for each compound following the extraction procedure described above. The R^2^ values for each standard curve exceeded 0.99.

### Rhodamine-Phalloidin staining of actin *in vivo*

A mid log culture of *S. cerevisiae* (diploid; BY4743) was diluted into fresh YPD to obtain 0.095 OD_600_/ml (~1−1.5×10^6^ cells/ml) and occidiofungin (1μg/ml; 0.5X MIC) or an equal volume of DMSO was added. Cultures were placed at 30°C with shaking for 0.5 or 2 hours. Formaldehyde (3.7% final) was added directly to the culture and cells incubated for 1.5 hours at room temperature. Formaldehyde was removed by vacuum filtration through 0.2μm nitrocellulose followed by several PBS washes. Cells were resuspended in PBS and permeabilized with the addition of an equal volume of 0.2% Triton X-100 PBS solution. Phalloidin-Tetramethylrhodamine B isothiocyanate (3.3mM in 100% DMSO; Sigma) was added to 6.6μM and cells stained for 30 minutes at room temperature. Unbound phalloidin-TRITC was removed by centrifugation at 8,000 *g* for 8 minutes at 20°C, the cells resuspended in PBS and added to concanavalinA treated glass slide and overlaid with VectaShield plus DAPI. Images were viewed using a Nikon 50i fluorescence microscope with a 100× oil immersion objective and Texas-Red and DAPI filter sets. Random images were captured using a Retiga EXi Black and White CCD Camera and Image Q software. All images were captured using the same exposure settings with image contrast altered post capture using CorelDraw.

### Efficacy studies using occidiofungin

The murine model of vulvovaginal candidiasis has been widely reported[77]. A variation of the method described was followed. Three groups of six mice were used to evaluate two concentrations of occidiofungin (100 μg and 50 μg) and vehicle control (0.3% Noble agar). Briefly, six to eight-week-old BALB/c mice were treated subcutaneously with 200 ng per mouse of β-Estradiol 17-valerate three days prior to inoculation with *C. albicans* (D-3). A subcutaneous dose of estradiol was administered every three days (D0, D3) until the end of the experiment to induce pseudo-estrus. Approximately a 20 μL intravaginal inoculation of a 2.5×10^6^ colony forming units (CFU)/mL of *C. albicans* defines day zero (D0) of the VVC study. On the same day of inoculation (D0), another subcutaneous injection of estradiol was made. Lyophilized powder of occidiofungin containing either 100 μg or 50 μg of occidiofungin was suspended in 20 μL of warm 0.3% Noble agar before intravaginal inoculation. Drug treatment was done on day 2 (D2), day 3 (D3) and day 4 (D4) of the study. On day 5 (D5), the vaginal lumen was lavaged with 100 μL of sterile PBS with a 200 μL pipette tip. Serial dilutions and total colony forming units per vaginal lavage were determined by plating on YPD plates containing 50 μg/mL of chloramphenicol. The colony forming units (CFUs) obtained from each lavage were counted on plates containing 30-300 colonies for determining the CFU/mL estimates. Body weight, signs of vaginal irritation such as swelling or bleeding and clinical signs of discomfort (stereotypical stretching behavior) were monitored. Statistical analyses (T-test) were done to compare the control group to treated groups and to compare differences between treated groups. All the analyses were 2-sided, with P < .05 considered statistically significant.

### Ethics Statement

Research Compliance’s Animal Welfare Office (AWO) supports Texas A&M’s Institutional Animal Care and Use Committees (IACUC), through which all faculty, staff, and students using animals, regardless of location or funding, must obtain approval before activities begin. The committee approved the study titled “Determination of efficacy of occidiofungin in the treatment of vulvo-vaginal candidiasis (IACUC number: IACUC 2017-0164)”. The specific national guidelines followed by Texas A&M’s AAALAC accredited animal facilities are the USDA animal welfare assurance regulations (Texas A&M registration: 74-R0012) and PHS NIH Guidelines (Texas A&M registration: A3893-01).

## Acknowledgements

We would like to acknowledge Joseph Sorg and Kathryn Ryan in the Biology Department at Texas A&M University for their erudite discussions. We would also like to thank Lawrence Dangott for his assistance with the pulldown assays. We also thank Dr. Susan Forsburg for providing the *Schizosaccharomyces pombe* 972h-strain.

## Supporting Information

S1 Table: Activity of occidiofungin against filamentous and non-filamentous fungi.

S2 Table: Occidiofungin exposure results in nuclear segregation defects in mitotic cultures of *S. cerevisiae* and *C. albicans*. Nuclear DNA was scored by DAPI staining of fixed cells treated with 0.5X MIC occidiofungin for 0.5, 1, and 2 hours at 30°C. Cells were binned into one of four categories based on bud morphology and DNA localization and the percentage of cells in each category are reported. Approximately 200 cells were scored for each time point and data from two separate experiments are shown.

S3 Table: Activity of occidiofungin against *S. cerevisiae* mutants deleted for genes linked to actin polymerization and depolymerization.

S4 Table: Activity of alkyne-OF compared to native occidiofungin

S5 Table: List of proteins pulled down exclusively by alkyne-OF using the affinity purification. Proteins in the cells highlighted in green are those that were found in the pulldown assays in both *S.pombe* and *S.cerevisiae*. Proteins in the cells that are not highlighted were found in the *S.pombe* assays only.

S1 Figure: Scheme of chemical addition of alkyne group to occidiofungin B and mass determination of alkyne-OF B.

S2 Figure: NMR data for Alkyne-occidiofungin A-D (^1^H 600 MHz and ^13^C 150 MHz in DMSO-d6) A) ^1^H NMR, B) COSY 2D NMR, C) TOCSY 2D NMR, D) HSQC 2D NMR, E) HMBC 2D NMR, and F) NOESY 2D NMR.

S3 Figure: Assignment of alkyne subunit and revised structure of diaminobutyric acid in the cyclic peptide. A) Expanded 1H NMR, B) Expanded TOCSY, C) Expanded HSQC, D) Expanded HMBC, E) Further expansion of HMBC, F) Expanded COSY NMR, G) Expanded TOCSY NMR, and H) Carbon and proton assignments of alkyne subunit.

S4 Figure: Induction of apoptosis by alkyne-OF: The ‘DMSO’ and ‘H_2_O_2_’ columns represent the negative and positive controls, respectively. The ‘WT’ column corresponds to cells treated with 1× MIC quantity of native occidiofungin and the last two panels represent treatment of cells with alkyne-OF at the concentration indicated. A) Externalization of phosphatidylserine demonstrated by the fluorescence of Annexin-V-Fluorescein, B) Release of reactive oxygen species indicated by the formation of rhodamine from dihydrorhodamine 123 and C) Double stranded breaks visualized by TUNEL assay, following treatment with native and alkyne-OF.

S5 Figure: Effect of occidiofungin on actin (a) polymerization and (b) depolymerization *in vitro*. Symbols are as follows: 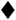- G-buffer (control), 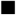 - G-buffer and pyrene actin, 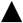 - Test buffer (1.5% β-cyclodextrin in PBS) and pyrene actin (control), X - 20 μL of test buffer containing 20 μg of occidiofungin and pyrene actin.

S6 Figure: Visualization of actin filaments: a) Untreated F-actin filaments stained with phalloidin 670 dye; Alkyne-OF treated F-actin filaments stained with azide derivatized AlexaFluor488 [(b)-(40x); (c)-(100x)]

